# Electrodeformation of White Blood Cells Enriched with Gold Nanoparticles

**DOI:** 10.1101/2021.09.25.461820

**Authors:** N. G. Hallfors, J. M. Teo, P. Bertone, C. Joshi, A. Orozaliev, M. N. Martin, A. F. Isakovic

## Abstract

The elasticity of white blood cells (WBCs) provides valuable insight into the condition of the cells themselves, the presence of some diseases, as well as immune system activity. In this work, we describe a novel process of refined control of WBCs’ elasticity through a combined use of gold nanoparticles (AuNPs) and the microelectrode array device. The capture and controlled deformation of gold nanoparticles enriched white blood cells *in vitro* are demonstrated and quantified. Gold nanoparticles enhance the effect of electrically induced deformation and make the DEP related processes more controllable.

## Introduction

Cells are subjected to a variety of mechanical forces in vivo, and the way they deform in response to mechanical, electrical, and biochemical stimuli relies on a combination of passive and active processes [1]. Red blood cells significantly deform as they travel throughout the body’s cappilary networks, which are at times smaller than the cells’ resting size. Diseases such as malaria and sickle cell anemia are associated with disruption of the cell membrane elasticity, leading to capillary blockages and a loss of oxygenation [2]. The deformability of cells has even been linked to cancer, where highly metastatic cells have been shown to be soft and deformable, allowing them to migrate through tissue into the blood stream [3, 4, 5].

In white blood cells (WBC), quantification of a cell’s elastic modulus via deformability measurements could provide insights into the physiological state of the cell. HL60 cells can differentiate into monocytes, granulocytes or macrophages, and mechanical deformation alone can distinguish which pathway the HL60 cell will take [6]. Neutrophil activation leads to reduced deformabillity, which has been demonstrated by morpho-rheological (MORE) analysis [7]. Monocytes from individuals afflicted by (Respiratory Tract Infection) RTI or (Acute Lung Injury) ALI both increased in size with staphylococcus stimulation, but only viral RTI monocytes displayed any measurable increase in deformation. These results indicate that size and deformation studies may be able to identify the presence of viral, bacterial, or other inflammatory diseases through lymphocyte mechanical analysis, implying a need for fast, reliable methods to measure mechanical properties.

A number of methods currently exist to deform and measure the elasticity of cells. Direct methods such as AFM and parallel plate rheometry [3,8,9] involve physically deforming a cell by applying force with a contact probe and measuring probe deflection. Optical tweezers and optical stretchers utilize laser light to deform cells and measure elastic modulus [5,10,11,12]. Optical deformation has even been used to demonstrate the identification of noncancerous, cancerous and metastatic cells in a mixed population [13].

Dielectrophoresis (DEP) is a method of manipulating particles suspended in a fluid medium, whereby a particle placed in a non-uniform electric field experiences a force through its dielectric response. The amplitude of the DEP forces experienced by the cells is modulated by the dielectric properties of the cell and surrounding media, and is expressed by Clausius– Mossotti (CM) factor, which will vary from -1 for a strongly repellent force, to +1 for a strongly attractive force [14]. Under certain conditions, a cell may become trapped and deformed by DEP, a phenomenon hereby referred to as electrodeformation.

A number of studies have utilized electrodeformation to controllably deform cells to measure mechanical characteristics. Electrodeformation of red blood cells (RBCs) have been performed with various devices utilizing DEP [15,16,17 and references therein], as well as more sophisticated trap and release microarrays for high throughput imaging and characterization [18]. The majority of such studies have involved red blood cells [19] due to their readily available nature and easily observable electrodeformation behavior, that follows the analytical prediction from the model to be discussed hereafter. We wish to emphasize that the overall review of the electrodeformation of RBCs is beyond the scope of this manuscript.

In contrast with RBCs, and despite a number of promising initial studies, a fast, reliable, highly parallelized, and scalable method for the controlled deformation and observation of white blood cells (WBCs) is yet to be realized. In the current work, a microelectrode array was designed to accommodate several cells at once, for rapid, parallel capture, deformation and imaging (Figures 1, 2 here)) of WBCs. Additional specific contribution of this report is the use of gold nanoparticles (AuNP), introduced to the cells to enhance the effect of electrodeformation.

**Figure 1.**
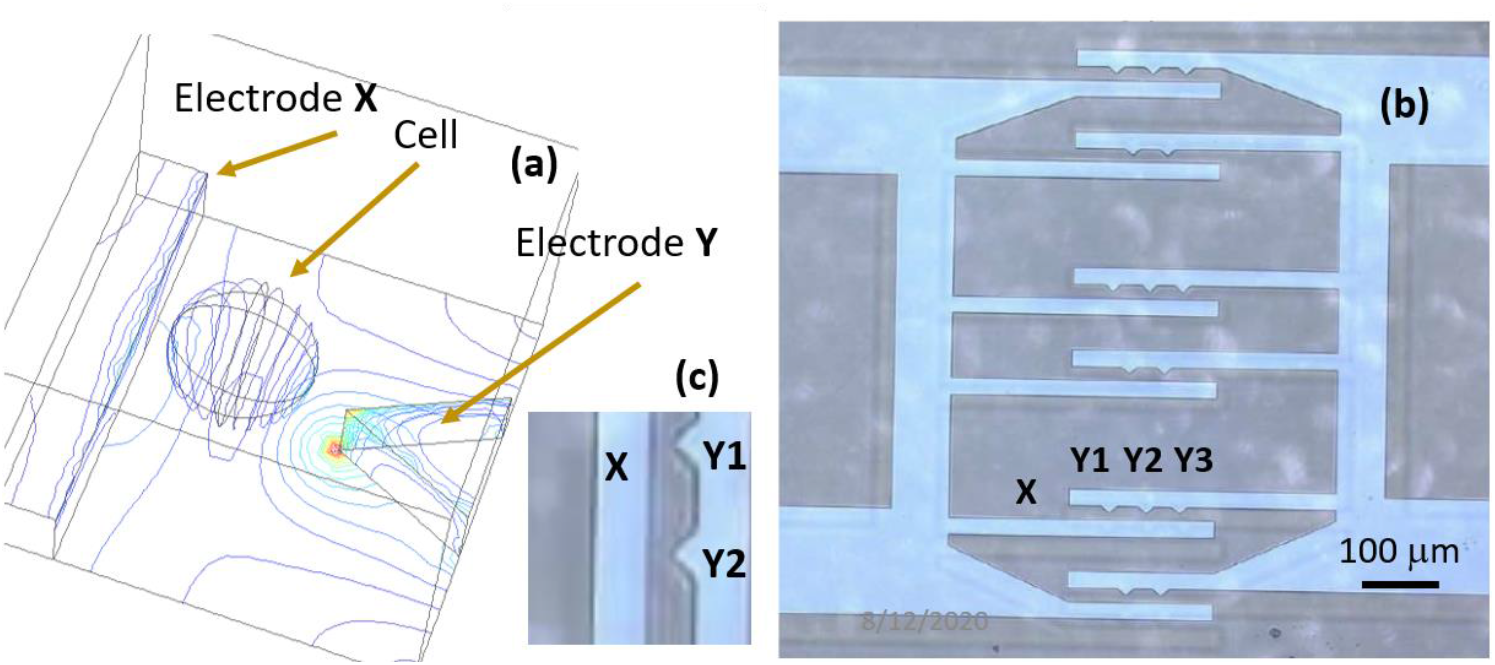
a) 3D numerical simulation of electrodes and cell (electrode height not to scale) showing electrical potential lines in a representative medium. b) Optical microscope image of ITO electrodes on glass. c) closeup of a fraction of electrode array at the region of highest field strength. Cells are attracted to this region by DEP forces, and fixed at electrode tips, where they are subsequently deformed.

**Figure 2.**
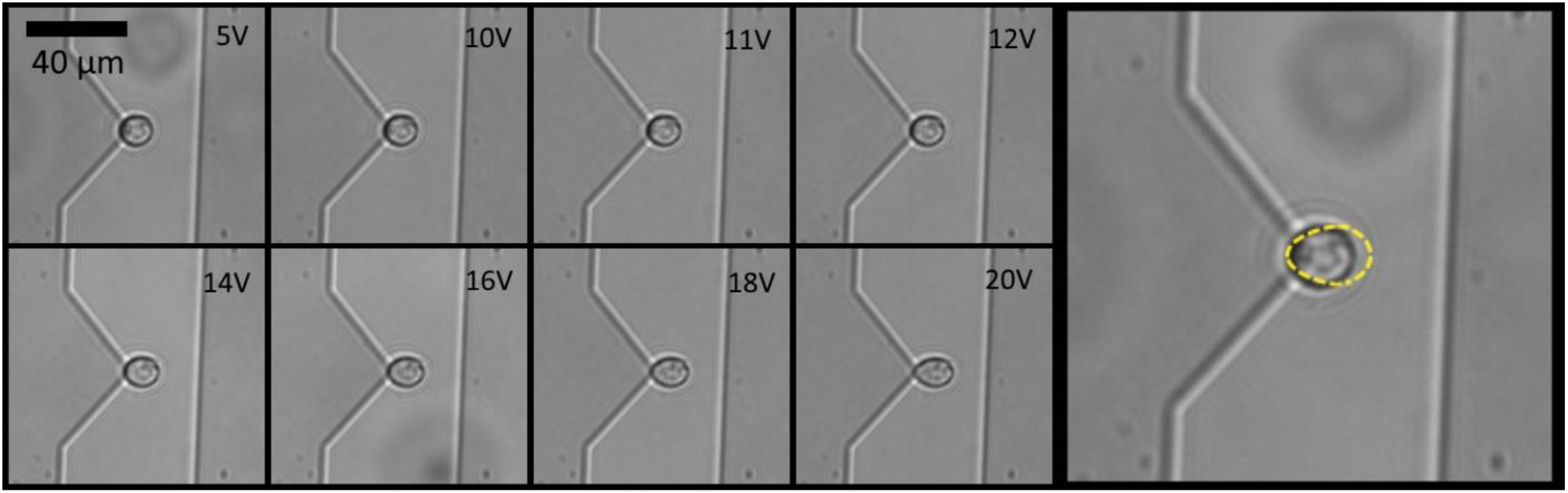
THP-1 Cell captured at electrode tip by DEP forces, and subsequently deformed. Right side shows an overlay of the 20V deformation on top of the resting cell. Supporting Materials contain the movie file of this process.

## METHODS

### Dielectrophoretic Electrodes on a Chip

Standard microfabrication techniques were used to produce indium tin oxide metal electrodes on a glass substrate. 150 nm thick indium tin oxide (ITO) was deposited by electron beam evaporation on a borosilicate glass wafer. A lift-off process was used to pattern electrodes on the substrate. The patterned electrodes were subjected to an annealing process at 400 deg C in air. The high temperature annealing of ITO while increasing oxidation results in an ITO film that is highly transparent and conductive (Figure 1, 2) [20,21,22]. Following lift off, glass wafers were diced with a dicing saw. More details are avialble in Supporting Online Materials.

### Cells

Human THP-1 and Jurkat cell lines (ATCC) were grown in RPMI 1640 Medium containing D-Glucose, HEPES, L-Glutamine, Sodium Bicarbonate, and Sodium Pyruvate (Gibco) supplemented with 10% Fetal Bovine Serum (FBS, Gibco) and 1% penicillin-streptomycin (Biosera). For cells with gold nanoparticles, cells were incubated in media with a small volume of added AuNP colloid for 24 hours [23]. Uptake of AuNP was verified visually and by X-ray fluorescence (XRF) (Figure 3, here).

**Figure 3.**
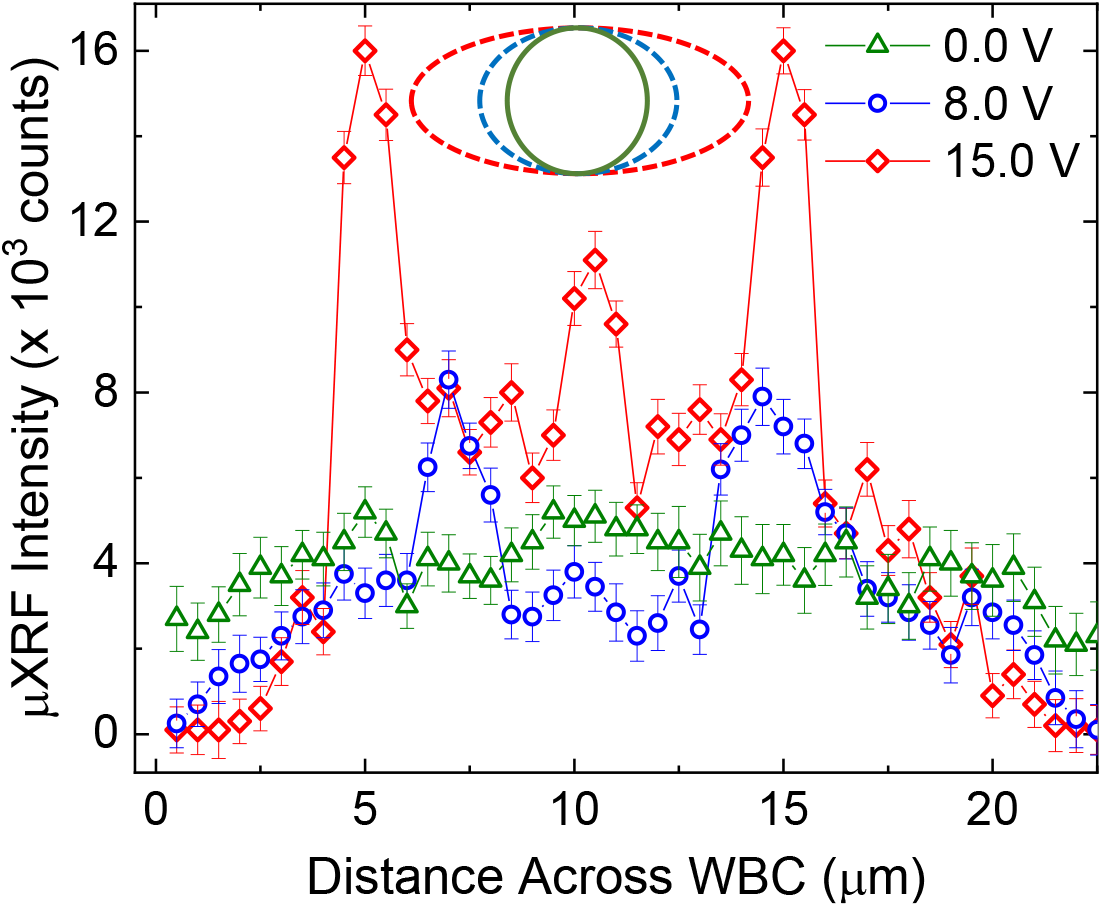
micro-X-ray fluorescence scans performed while cell and gold nanoparticles are under the influence of applied electric field. Zero voltage shows relatively evenly distributed Au NPs. As the voltage is increased, the cell is deformed, and increased concentration of Au NPs in the membrane region. The upper centered insert shows sketched deformation of the WBC (not to scale), under the assumption that the deformation is elliptical.

### Gold Nanoparticles (AuNP)

Synthesis of naked AuNPs was performed following the method of Martin *et al*. [24]. AuNP are generally considered biocompatible and are being investigated for their applications in medicine and research as carriers for bioactive compounds [23] as contrast agents [24] or as radiation absorbent materials [25]. A 40 mL capacity clean borosilicate glass vial containing 9.25 g of de-ionized water was mixed with an aqueous gold precursor solution 0.1 g (∼100 μL) containing 50 mM HAuCl_4_/HCl producing a light yellow solution. To this light-yellow solution 0.65 g (∼650 μL) of freshly prepared aqueous 50 mM NaBH_4_/NaOH was added rapidly while vortexing. Upon the addition of the alkaline borohydride solution the reaction mixture immediately turned red, signaling the nucleation of gold nanoparticles at room temperature, and was vortexed for one minute. The ruby red AuNP solution was then placed in a metal heating block (already at 250 °C) for three minutes to grow the AuNPs and improve the monodispersity. The vial was then quenched for 30 seconds under running water to arrest the kinetic growth of NPs. These hydroxide-stabilized naked-AuNPs were then used for the WBC experiments here. More details in Supporting Online Materials.

### Apparatus

Electrode array chips were placed on glass slides and connected to a Rigol DG4062 function generator via copper tape and coaxial cable. Cells were centrifuged at 1000 rpm for 5 minutes, and RPMI was aspirated. A DEP media composed of 8.5% (w/v) sucrose and 0.3% (w/v) dextrose in distilled water was adjusted to 10 mS/m by adding a small volume of Phosphate buffered saline (PBS). Cells were resuspended in the DEP media at low density. A small volume of cell suspension was placed on the electrodes, and AC current was applied. For deformation experiments, frequency was fixed at 700 MHz, while voltage was stepped from 0 – 20 Vpp in 1V increments. Cells were imaged on a Zeiss Observer Z1 microscope in bright field mode.

DEP forces induced by the field asymmetry pulled cells toward the electrode tips, where field was highest (Fig. 2). Once at the tip, cells become fixed in place and began to deform as voltage was increased (Fig. 2). Voltages above 20.0 V generally caused cells to span the electrode gap, doing permanent damage to the cell, so voltage was limited to 20.0 V for the purpose of this study. The electrically induced deformation is reversable and repeatable for at least 10 cycles under the conditions outlined in this Report.

### Image analysis

Images were imported to ImageJ, and cell dimensions were manually measured, frame by frame. An elliptical selector was used to best fit the cell during deformation, and all cells are assumed to be elliptical at all time points. Ellipse geometries were then exported for analysis.

## RESULTS

### Electrode Array Fabrication

Indium tin oxide on glass was chosen for electrode fabrication due to ideal optical and electrical properties. The electrode geometry was designed in such a way as to accommodate single cells, positioned by DEP forces prior to deformation (**Figure 1**). The small gap size allows us to keep voltage low, while the electrode spacing was chosen to balance the need for separation between cells with parallelization of the process. Additional details of the microfabrication process are availabel in the Supporting Online Materials file.

### X-ray microfluorescence for detection of gold nanoparticles absorption

Our early tests demosntrated promising response of white blood cells to DEP forces in the presence of gold nanoparticles (Au NPs). This has motivated the desire to establish where exactly are Au NPs located under the experimental conditions in this report.

To this end, electrodeformation experiments were conducted while electrode array device and WBCs were under X-ray illumination. Figure 3 here shows the result of changing spatial concentration of Au NPs as scanning with a sub-micron resolution (nominal resolution 250 nm +/- 75 nm) is performed. Prior to the application of electric field the spatial distribution of Au NPs is apprixmately even, as there is no region inside or outside the WBCs that generates stronger fluorescent signal. The situation changes appreciably after the electric filed is turned on, so at 8.0 V we see the peaks the separation of which roughly approximates the width of the white blood cells tudied. It is also possible to detect some decrease in the fluorescence signal outside the cell and at the cell’s center. Further increase of the applied electric field produces additional peaks in the fluorescence distribution, indicating that a majority of Au NPs become adsorbed at the cell membrane (either inside or outside the cell), with some additional concentration in the area of cell’s center. We note that there are uncertainties in these data originating in the averaging of the XRF signal over several cells and in the positioning. The uncertainties for the horizontal axis (distance across) measurement are ommitted, as they are well approximated by the size of the symbol. This approach could be seen as a pilot study leading into additional insights about function of the cells when enrpiched with AuNPs [26, 27], but we emphasize the result presented in Figure 3 serves only to confirm that there is a change in the spatial distribution of AuNPs in and around the WBCs as one changes the applied electric field.

### Frequency dependent response

DEP forces exerted upon a cell are dependent upon frequency, which is often reflected in the behaviour of the CM factor. The CM factor profile is different for each cell type, so understanding the CM profile is an important piece of any DEP study. A study of the frequency response was performed on both THP-1 and Jurkat’s cells. This was accomplished by keeping the voltage constant at 10V while varying the signal frequency and measuring the time needed for nearby cells to reach the electrode tip. This time is inversely related to the attractive force, and serves as a good initial approximation of the CM factor [28, 29]. Due to the design of our device, negative DEP forces are very small, and this method fails as the frequency approaches the cell’s crossover value and time goes to “infinity”.

Extrapolation is used to predict the cell’s crossover value based on the measurements approaching negative DEP (Figure 4 (L)). Specifically, polynomial fitting and extrapoloation lead us to values for crossover frrequencies *f*_xo1,Jurkat_ = 21.5 kHz +/- 3.4 kHz and *f*_xo1,THP-1_ = 57.7 kHz +/- 3.4 kHz. These values are broadly in agreement with a number of related studies of white blood cells [30-34]. We wish to emphasize that the main purpose of this report is the is the deformability study, so, this measurement has been conducted primarily as an overall quality check, and its focus was not to suggest significantly different values of crossover frequencies for Jurkat and THP-1 white blood cells are called for. These experimental results motivated our need for better understanding the effect the introduction of AuNPs into WBCs on electrostatic properties of WBCs. With this in mind, we modeled dielectric response of WBCs using Claussius-Mossotti approach in a double-shell approximation (See Supporting Materials).

**Fig. 4.**
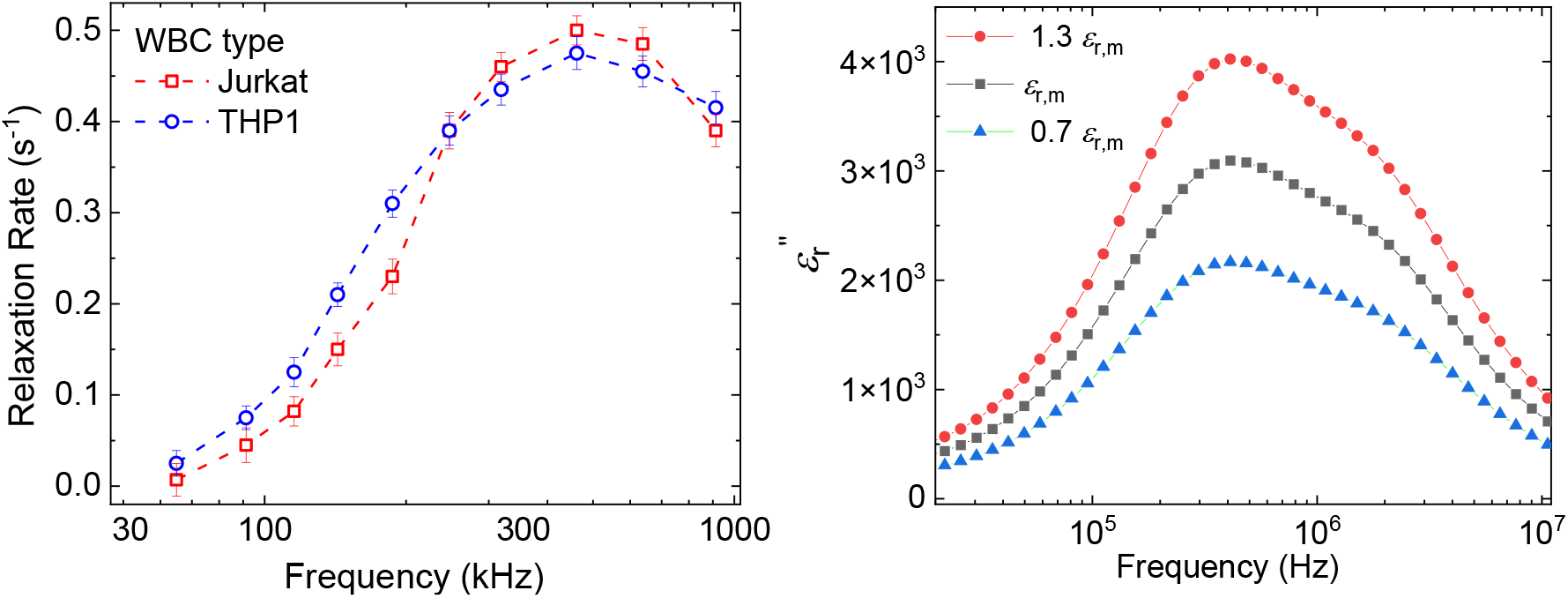
(L) The inverse of the time of the “attraction until rest” is recorded at different frequencies and the fixed applied electric field. In the text we elaborate how these data are (1) fit to a polynomial and then (2) extrapolated to obtain the reported crossover frequencies. (R) Claussius-Mossotti multi-parameter model indicates a possible range of changes of the relative impedance upon addition of Au NPs that allows WBCs to achieve the peak at a similar frequency observed in (L).

We have used output of this model and the prescription offered by [29] to generate the plot in the Fig. 4. (R), for the imaginary component of complex permitivity. The white blood cell parameters are available (references here and Supporting Materials), and approximating the influence of AuNPs as a change of the relativity permitivity of the medium WBCs are used in for this experiment. Understandably, this approxmation is a simplification, but, we suggest that, to a first order, scalar modification of permitiy is one way to understand the role of AuNPs. Details of the model are provided in Supporting Materials.

### Deformation study

Two cell lines were studied: THP-1 monocytes and Jurkat’s T-lymphocytes. The cells were deformed without gold nanoparticles (AuNPs) modification, then after incubation with plain AuNPs, and AuNPs modified with PEG and citrate. Cells were suspended in isotonic, nonconducting media for DEP. DEP forces attracted single cells to the electrode tip, where they were trapped and subsequently deformed. A gradual deformation was induced by increasing voltage in 1.0 Volts increments at 700kHz. Very little deformation is seen until the voltage reaches a value of 5.0 Volts for most cells, after which cells are stretched into a shape well approximated by an ellipsoid (ellipse in 2D), with the aspect ratio of the ellipse increasing as the applied voltage grows. Voltage was limited to a 20V maximum, as beyond this point cells suffered permanent damage. This is not an impediment to scale-up of this method, as we will see that the voltages below 20.0 V are sufficient for controllable deformation and cell sorting protocols. If cell density is too high, the likelihood of multiple cells chaining together to span the electrode gap increases. In this case, the cells do not fully deform, but become permanently trapped at the electrodes, so we have avoided performing measurements for more than one cell. It is not impossible to rely on multi-cell capture, but the findings from such work require more detailed statistical analysis, and this report focuses on simple, easy-to-repeat experiments.

Figure 5 shows raw deformation data for both types of cells and for variations in the utilization of AuNPs (PEG, citrate). Some observations that follow from this are:

**Fig. 5.**
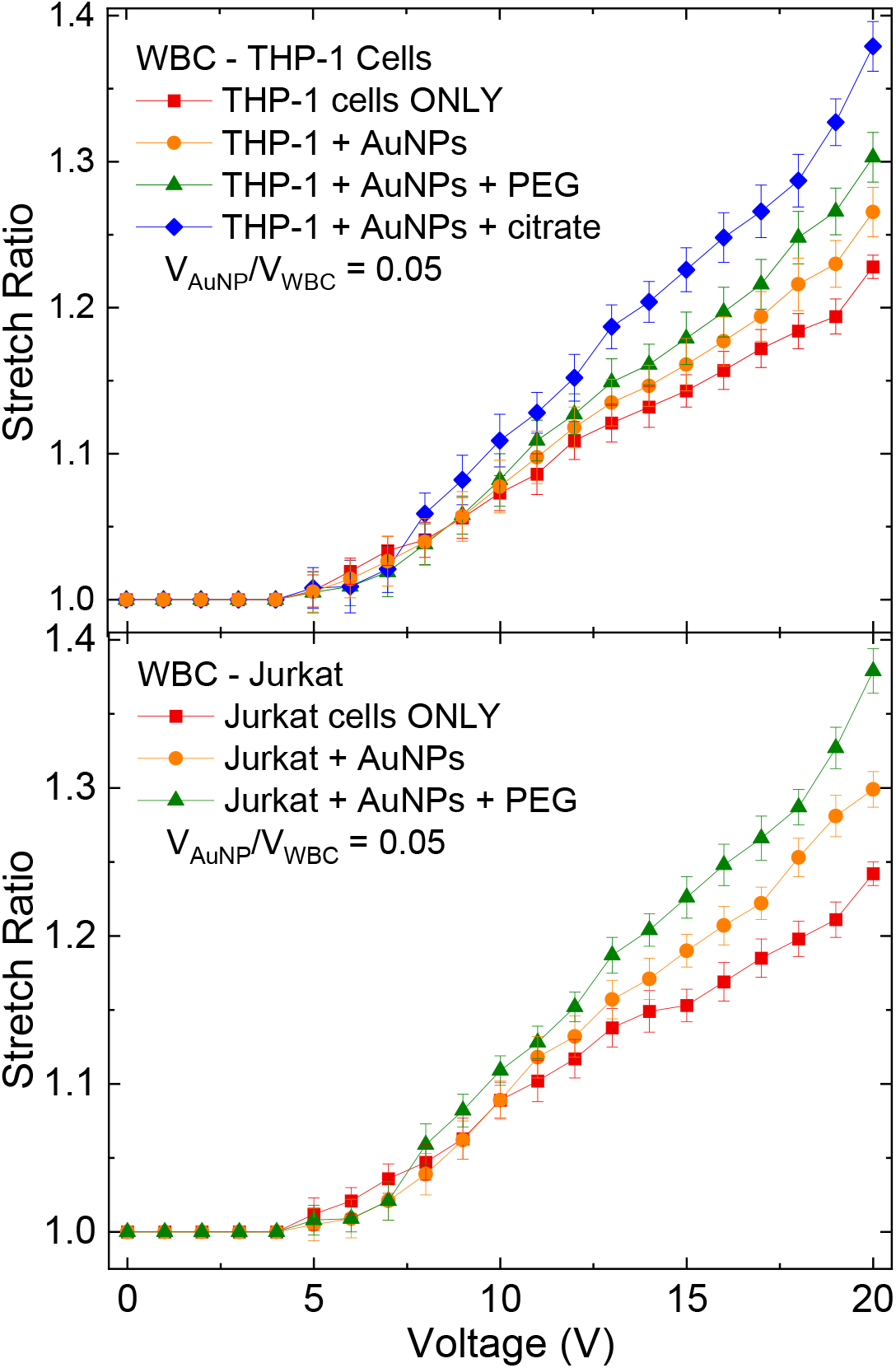
(top) Stretch ratio measured as a function of the applied voltage for THP-1 white blood cells (WBC). In addition to “cells only” and “nanoparticles enriched WBCs”, we used PEG-ylated and citrated solutions of Au NPs, as discussed in the main text. (bottom) same study, but for the Jurkat type WBCs. Elliptical deformation is assumed during analysis of cells’ images.

a. It is apparent that very little deformation occurs below 5.0 V
b. It takes some additional voltage increase (+2 to 3 Volts, in general) to start observing statistically significant differences for various modifications of WBC with AuNPs.
c. As expected, the larger is the applied voltage, the more difference between deformation curves is observed
d. The deformation curves, the stretch ratio vs the applied voltage, while generally non-linear, are not as apparently quadratic in voltage, as one would expect from numerous electrodeformability studies conducted for red blood cells.

In order to better understand data in Fig. 5, and ultimately to check for the predicted quadratic dependence [14, 27] of the stretch ratio on applied voltage, we have performed the fit of the stretch ratio as a linear function of the square of the applied voltage. From the deformation data, we are able to calculate the shear elastic modulus for the cells by a modified Maxwell force model from Engelhardt and Sackmann [17].

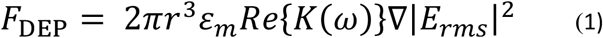

where *r* is the particle radius, *ε*_*m*_ is the absolute permitivity of the fluid, and

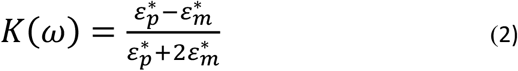

is the Clausius–Mossotti factor, which describes the polarizability of the particle. *E*_*rms*_ is the root mean square amplitude of the electric field. These calculations rely on the Maxwell stress, which predicts a dependence of the deformation as the square of the voltage.

As can be seen from Figure 6 here, the data demonstrate an overall good fit to V^2^ (*κ*_R_^2^ in the range 0.95 and 0.98 for all fitting shown), but with a caveat that there is a region of data overperforming and underperforming the parabolic fit. One way to interpret data in Figures 5 and 6 here is to propose the existence of different regimes of electrodeformability for white blood cells. For example, based on the intersection points between the data and the straight fit lines in Fig. 6 it seems that these regimes would be, approximately (i) 0.0 V – 8.0 V, (ii) 8.0 V – 15.0 V, and (iii) 15.0 V – 20.0 V. Assuming this empirically driven qualitative argument has a basis in the fundamental response of {WBCs + AuNPs} system, it would indicate that a more detailed, molecular level model of the behavior of {WBCs + AuNPs} system in electric field is called for. Development of such model is beyond the scope of this report, however. It would likely start by accounting for (i) an increase in dielectric constant due to the insertion of AuNPs and/or (ii) charge separation of the counterions from the double layer occurring before migration of AuNPs.

**Figure 6.**
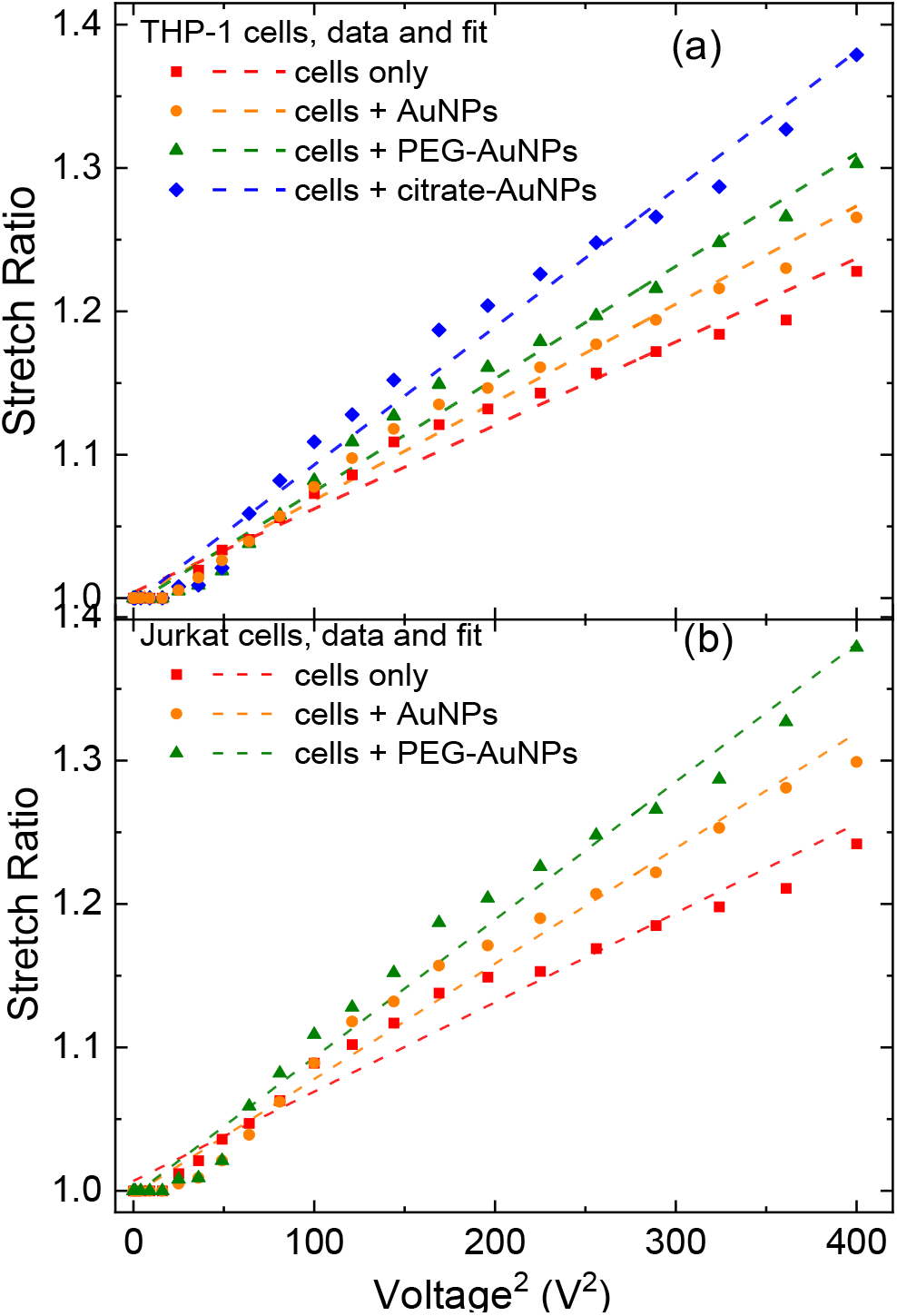
(a) Stretch ratio plotted as a function of the squared applied voltage, for THP-1 white blood cells (WBC), for the purpose of linear fit, Points represent the same values as in Fig. 5, while the dashed line represent the fit values. It is clear that there are different deformation regimes, as indicated by the crossover points (where the imaginary line connecting the points crosses the line of the fit); (b) same analysis for Jurkat cells.

Finally, based on electrodeformability data from Fig. 5, and fitting analysis in Fig. 6, we have determined the elastic modulus for the types of WBC we studied here. Results are reported in Fig. 7 show a significant reduction in the effective elastic modulus for both THP-1 and Jurkat cells with the addition of AuNP, and further reduction with nanoparticle modification. Some possible mechanisms for AuNP effected modulus changes are:

**Figure 7.**
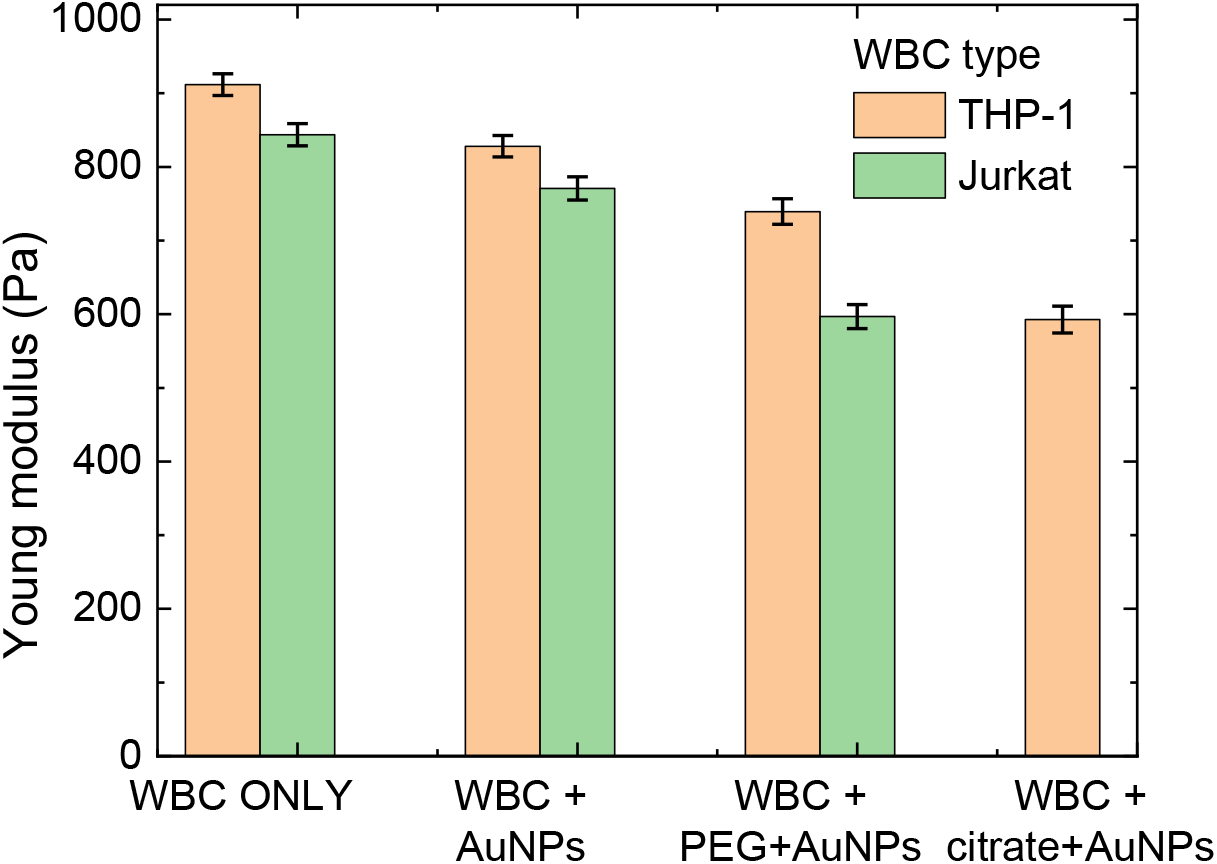
Values of the Young modulus for different experimental cases in this Report.

1. Particles embedded within the cell membrane or floating within the cytoplasm change the electrical impedance enough to affect the permittivity of the system.
2. Nanoparticles provide additional active surface area for electrical forces, effectively increasing the surface area of the cell without increasing its resting radius.
3. Furthermore, AuNP modified with molecular conjugates PEG and citrate showed additional changes to the effective elastic modulus for WBCs.

## Conclusions

We have fabricated a novel microelectrode array for the controlled deformation of white blood cells. THP-1 monocytes and Jurkat’s T-lymphocytes were enriched with gold nanoparticles, deformed with electrostatic forces, and based on the geometry of the device, applied voltage and cell size, Maxwell tension and cell elastic modulus were determined.

This method was developed because, despite the existence of several methods for measuring white blood cells’ elastic moduli, they often give different results for similar cell types. Introduction of AuNPs helps in this regard. In addition to cell elastic modulus, we report on spatial resolution of the adsorbed AuNPs, and on crossover frequencies in the standard Claussius-Mossotti model of response.

The nature of the electrodeformation data indicates that novel efforts are needed to better understand the response on white blood cells to electric fields on molecular level. The device we have developed is likely to lead to a highly parallel measurement of cells with a high degree of confidence for high throughput characterization. Overall, these results represent a useful addition to the rapidly growing field of on-chip quantifiable cells’ properties [35-48].

## Contributions

JT and AFI conceived the study. NGH led the effort in design and fabrication of devices with the help from AO and AFI. NGH and AFI performed measurements. Analysis was done by NGH, PB, and AFI. MNM and CS prepared AuNPs. The draft of the report was written by NGH, AFI, PB and all co-authors help edit it.

## Data availability

Data from this report are available upon reasonable request.

## Acknowledgments

The authors thank KUST BME Dept for the use of equipment, KUST – MASDAR Institute for the access to Microfabrication Facility NYU-Abu Dhabi for the assistance in mask making. We express our gratitude to B. Samara and L. George (KUST) for technical assistance. We are especially thankful to our colleagues at Brookhaven National Lab for the access to X-ray fluorescence setup and for providing additional cells and chemicals. BNL-CFN and BNL-NSLS-II are funded by US Department of Energy. NGH acknowledges Graduate Scholarship Fund at KUST. JT, MNM and AFI acknowledge KUST Internal Research Funds. AFI acknowledges hospitality and support at Cornell CNF, supported by US NSF.

## Supporting Online Materials

This report is accompanied by the Supporting Materials file which contains additional details on (i) properties of AuNPs, (ii) details of the microfabrication process, (iii) details of the modeling of Claussius-Mossotti {WBC + AuNPs} system.

## References

[1] Bongrand, P., Benoliel, A. M., & Richelme, F. (1997). Mechanical deformation of monocytic THP-1 cells: occurrence of two seqential phases with differential sensitivity to metabolic inhibitors. Experimental Biology Online-EBO, 2(5), 1–14. DOI: 10.1007/s00898-997-0005-8

[2] Ahmad, I. L., & Ahmad, M. R. (2014). Trends in characterizing single cell’s stiffness properties. Micro and Nano Systems Letters, 2(1), 8. DOI: https://doi.org/10.1186/s40486-014-0008-5

[3] Cross, S. E., Jin, Y. S., Rao, J., & Gimzewski, J. K. (2007). Nanomechanical analysis of cells from cancer patients. Nature Nanotechnology 2(12), 780. DOI: https://doi.org/10.1038/nnano.2007.388

[4] Baker, E. L., Lu, J., Yu, D., Bonnecaze, R. T., & Zaman, M. H. (2010). Cancer cell stiffness: integrated roles of three-dimensional matrix stiffness and transforming potential. Biophysical Journal, 99(7), 2048–2057. DOI: 10.1016/j.bpj.2010.07.051

[5] Guck, J., Schinkinger, S., Lincoln, B., Wottawah, F., Ebert, S., Romeyke, M., … & Käs, J. (2005). Optical deformability as an inherent cell marker for testing malignant transformation and metastatic competence. Biophysical Journal, 88(5), 3689–3698. DOI: https://doi.org/10.1529/biophysj.104.045476

[6] Otto, O., Rosendahl, P., Mietke, A., Golfier, S., Herold, C., Klaue, D., … & Wobus, M. (2015). Real-time deformability cytometry: on-the-fly cell mechanical phenotyping. Nature Methods, 12(3), 199. DOI: 10.1038/nmeth.3281

[7] Toepfner, N., Herold, C., Otto, O., Rosendahl, P., Jacobi, A., Kräter, M., … & Ranford-Cartwright, L. (2018). Detection of human disease conditions by single-cell morpho-rheological phenotyping of blood. eLife 7, e29213. DOI: 10.7554/eLife.29213

[8] Radmacher, M., Fritz, M., Kacher, C. M., Cleveland, J. P., & Hansma, P. K. (1996). Measuring the viscoelastic properties of human platelets with the atomic force microscope. Biophysical Journal, 70(1), 556–567. https://doi.org/10.1016/S0006-3495(96)79602-9

[9] Wu, P. H., Aroush, D. R., Asnacios, A., Chen, W. C., Dokukin, M. E., Doss, B. L., Durand-Smet, P., Ekpenyong, A., Guck, J., Guz, N. V., Janmey, P. A., Lee, J., Moore, N. M., Ott, A., Poh, Y. C., Ros, R., Sander, M., Sokolov, I., Staunton, J. R., Wang, N., … Wirtz, D. (2018). A comparison of methods to assess cell mechanical properties. Nature Methods, 15(7), 491–498. DOI: 10.1038/s41592-018-0015-1

[10] Hwang, H., Choi, Y. J., Choi, W., Kim, S. H., Jang, J., & Park, J. K. (2008). Interactive manipulation of blood cells using a lens-integrated liquid crystal display based optoelectronic tweezers system. Electrophoresis, 29(6), 1203–1212. DOI:10.1002/elps.200700415

[11] Henon, S., Lenormand, G., Richert, A., & Gallet, F. (1999). A new determination of the shear modulus of the human erythrocyte membrane using optical tweezers. Biophysical journal, 76(2), 1145–1151. DOI: 10.1016/S0006-3495(99)77279-6

[12] Mills, J. P., Qie, L., Dao, M., Lim, C. T., & Suresh, S. (2004). Nonlinear elastic and viscoelastic deformation of the human red blood cell with optical tweezers. MCB-TECH SCIENCE PRESS-, 1, 169–180. DOI: doi:10.3970/mcb.2004.001.169

[13] Guck, J., Ananthakrishnan, R., Mahmood, H., Moon, T. J., Cunningham, C. C., & Käs, J. (2001). The optical stretcher: a novel laser tool to micromanipulate cells. Biophysical Journal, 81(2), 767–784. DOI: 10.1016/S0006-3495(01)75740-2

[14] Pethig, R. (2017). Dielectrophoresis: Theory, Methodology and Biological Applications. Wiley. ISBN 978-1118671450

[15] Guido, I., Jaeger, M. S., & Duschl, C. (2011). Dielectrophoretic stretching of cells allows for characterization of their mechanical properties. European Biophysics Journal, 40(3), 281288. DOI: 10.1007/s00249-010-0646-3

[16] Thom, F., & Gollek, H. (2006). Calculation of mechanical properties of human red cells based on electrically induced deformation experiments. Journal of Electrostatics, 64(1), 53 DOI: 10.1016/j.cryobiol.2009.04.001

[17] Engelhardt, H., & Sackmann, E. (1988). On the measurement of shear elastic moduli and viscosities of erythrocyte plasma membranes by transient deformation in high frequency electric fields. Biophysical Journal, 54(3), 495–508. DOI: 10.1016/S0006-3495(88)82982-5

[18] Doh, I., Lee, W. C., Cho, Y. H., Pisano, A. P., & Kuypers, F. A. (2012). Deformation measurement of individual cells in large populations using a single-cell microchamber array chip. Appl, Phys Lett. 100(17), 173702. DOI: 10.1063/1.4704923

[19] Du, E., Dao, M., & Suresh, S. (2014). Quantitative biomechanics of healthy and diseased human red blood cells using dielectrophoresis in a microfluidic system. Extreme Mechanics Letters 1, 35–41. DOI: 10.1016/j.eml.2014.11.006

[20] Wang, R. X., Beling, C. D., hng, S., Djurišić, A. B., Ling, C. C., Kwong, C., & Li, S. (2005). Influence of annealing temperature and environment on the properties of indium tin oxide thin films. Journal of Physics D: Applied Physics, 38(12), 2000. DOI: 10.1088/0022-3727/38/12/022

[21] Song, S., Yang, T., Liu, J., Xin, Y., Li, Y., & Han, S. (2011). Rapid thermal annealing of ITO films. Applied Surface Science, 257(16), 7061–7064. DOI: https://doi.org/10.1016/j.apsusc.2011.03.009

[22] Xu, Z., Chen, P., Wu, Z., Xu, F., Yang, G., Liu, B., … & Zheng, Y. (2014). Influence of thermal annealing on electrical and optical properties of indium tin oxide thin films. Materials Science in Semiconductor Processing 26, 588–592. DOI: 10.1016/j.mssp.2014.05.026

[23] Rana, S., Bajaj, A., Mout, R., Rotello, V., (2012) Monolayer coated gold nanoparticles for delivery applications, Advanced Drug Delivery Reviews, 64(2), 200–216, DOI: https://doi.org/10.1016/j.addr.2011.08.006

[23] C. Frede, I. Fortunati, V. Weber, N. Rossetto, F. Bertasi, L. Petrelli, et al., “Evaluation of gold nanoparticles toxicity towards human endothelial cells under static and flow conditions,” Microvascular Research, 97, pp. 147–155, 2015.

[24] Cheheltani, R., Ezzibdeh, R. M., Chhour, P., Pulaparthi, K., Kim, J., Jurcova, M., Hsu, J. C., Blundell, C., Litt, H. I., Ferrari, V. A., Allcock, H. R., Sehgal, C. M., & Cormode, D. P. (2016). Tunable, biodegradable gold nanoparticles as contrast agents for computed tomography and photoacoustic imaging. Biomaterials 102, 87–97. DOI: https://doi.org/10.1016/j.biomaterials.2016.06.015

[25] Rahman, W.N., Bishara, N., Ackerly, T. He, et al. (2019) Enhancement of radiation effects by gold nanoparticles for superficial radiation therapy, Nanomedicine: 5(2), 136 https://doi.org/10.1016/j.nano.2009.01.014.

[26] L. E. Cole, R. D. Ross, J. M. R. Tilley, T Vargo-Gogola, R. K. Roeder, Nanomedicine 10, 321 (2015). DOI: 10.2217/NNM.14.171

[27] T. Sasaya, N. Sunaguchi, K. Hyodo, T. Zeniya, T. Yuasa, Scientific Reports 7, 5742 (2017), DOI:10.1038/s41598-017-05179-2

[26] Gierhart, B, Howitt, D, Chen, S, Smith, R and Collins, S. D. (2007) Frequency Dependence of Gold Nanoparticle Super assembly by Dielectrophoresis, Langmuir, 23(24), 12450–12456 DOI: https://doi.org10.1021/la701472y

[27] H. A. Pohl and H. Pohl, Dielectrophoresis: the behavior of neutral matter in nonuniform electric fields vol. 80: Cambridge university press Cambridge, 1978. DOI: https://doi.org/10.1086/411635

[28] M. C. Purnell, M. B. A. Butawan, R. D. Ramsey, Physiol. Reports 6, e13722 (2018). https://doi.org/10.14814/phy2.13722

[29] C. S. Sodhi, L. C. de Sena Monterio OZelim, P. N Rathie, Heliyon 7, e06606 (2021). https://doi.org/10.1016/j.heliyon.2021.e06606

[30] Gabriel, S., Lau, R. W., Gabriel, C. The dielectric properties of biological tissues: II. Measurements in the frequency range 10 Hz to 20 GHz. Physics in Medicine and Biology, 41(11), 2251 (1996).

[31] J. Spadavecchia, D. Movia, C. Moore, C. M. Maguire, H. Moustaoui, S. Casale, Y. Volkov, A. Prina-Mello, Intl. J. Nanomed. 11, 791, (2016). https://doi.org/10.2147/IJN.S97476

[32] Jin, Q., Verdier, C., Singh, P., Aubry, N., Chotard-Ghodsnia, R., & Duperray, A. (2007). Migration and deformation of leukocytes in pressure driven flows. Mech. Res. Comms., 34(5-6), 411.

[33] Cemazar, J., Ghosh, A., & Davalos, R. V. (2017). Electrical manipulation and sorting of cells. In Microtechnology for Cell Manipulation and Sorting (pp. 57–92). Springer, Cham.

[34] R. K. Anand, E. S. Johnson, and D. T. Chiu, Negative Dielectrophoretic Capture and Repulsion of Single Cells at a Bipolar Electrode: The Impact of Faradaic Ion Enrichment and Depletion, J. Am. Chem. Soc. 137, 776, (2015). DOI: https://doi.org/10.1021/ja5102689

[35] Fuhr, G., Glasser, H., Müller, T., & Schnelle, T. (1994). Cell manipulation and cultivation under ac electric field influence in highly conductive culture media. Biochimica et Biophysica Acta (BBA)-General Subjects, 1201(3), 353. DOI: https://doi.org/10.1016/0304-4165(94)90062-0

[36] N. Piacentini, G. Mernier, R. Tornay, and P. Renaud, Separation of platelets from other blood cells in continuous-flow by dielectrophoresis field-flow-fractionation, Biomicrofluidics, 5, 034122, 2011. DOI: 10.1063/1.3640045

[37] D. Chen and H. Du, A dielectrophoretic barrier-based microsystem for separation of microparticles, Microfluidics and Nanofluidics, vol. 3, pp. 603–610, 2007. https://doi.org/10.1007/s10404-007-0151-x

[38] Morimoto, A., Mogami, T., Watanabe, M. Iijima, et al. (2015). High-density dielectrophoretic microwell array for detection, capture, and single-cell analysis of rare tumor cells in peripheral blood. PLoS One, 10(6), e0130418. DOI: 10.1371/journal.pone.0130418

[39] Borgatti, M., Altomare, L., Baruffa, M. Fabbri,, et al. (2005). Separation of white blood cells from erythrocytes on a dielectrophoresis (DEP) based ‘Lab-on-a-chip’ device. International journal of molecular medicine, 15(6), 913–920. DOI: https://pubmed.ncbi.nlm.nih.gov/15870893/

[40] Yang, J., Huang, Y., Wang, X. B., Becker, F. F., & Gascoyne, P. R. (2000). Differential analysis of human leukocytes by dielectrophoretic field-flow-fractionation. Biophysical journal, 78(5), 2680–2689. DOI: 10.1016/S0006-3495(00)76812-3

[41] Zheng, Y., Nguyen, J., Wei, Y., & Sun, Y. (2013). Recent advances in microfluidic techniques for single-cell biophysical characterization. Lab on a Chip, 13(13), 2464–2483. DOI: https://doi.org/10.1039/C3LC50355K

[42] Yang, L., Banada, P., Bhunia, A., and R. Bashir, Effects of Dielectrophoresis on Growth, Viability and Immuno-reactivity of Listeria monocytogenes. Journal of Biological Engineering, 2, 1, 2008. 134 DOI: 10.1186/1754-1611-2-6

[43] Pan, D. C., Myerson, J. W., Brenner, J. S., Patel, P. N., Anselmo, A. C., Mitragotri, S., & Muzykantov, V. (2018). Nanoparticle Properties Modulate Their Attachment and Effect on Carrier Red Blood Cells. Scientific Reports, 8(1).

[44] Zhang, H., Chang, H., & Neuzil, P. (2019). DEP-on-a-Chip: Dielectrophoresis Applied to Microfluidic Platforms. Micromachines, 10(6), 423.

[45] Chen, J., Abdelgawad, M., Yu, L., Shakiba, N., Chien, W.-Y., Lu, Z., … Sun, Y. (2011). Electrodeformation for single cell mechanical characterization. J. Micromech. Microeng., 21(5), 054012. doi: 10.1088/0960-1317/21/5/054012

[46] Bai, G., Li, Y., Chu, H. K., Wang, K., Tan, Q., Xiong, J., & Sun, D. (2017). Characterization of biomechanical properties of cells through dielectrophoresis-based cell stretching and actin cytoskeleton modeling. BioMedical Engineering OnLine, 16(1).

[47] J. K. Seo, T. K. Bera, H. Kwon, R. Sadlier, Comp. Math. Methods Medicine 2013, 353849 (2013). https://doi.org/10.1155/2013/353849

[48] T. N. G. Adams, K. M. Leonard, A. R. Minerick, Biomicrofluidics 7, 064114 (2013). http://dx.doi.org/10.1063/1.4833095

