## Supplemental Online Materials for "Electrodeformation of White Blood Cells Enriched with Gold Nanoparticles"

Supporting Online Materials for the manuscript "***Electrodeformation of White Blood Cells Enriched with Gold Nanoparticles***"

This manuscript brings together several (sub-)disciplines, so we feel the overall structure of the discussion and data in the main text is better understood if the information in this document is relied on, for one or more phases of work.

Each slide has a theme

- SOM-1 output of numerical modeling of the distribution of the electric field lines with the electrode geometry we relied on
- SOM-2 microfabrication process used to make the electrodes with some intermediate and final optical microscope images
- SOM-3 a movie depicting the deformation of a single white blood cell with gold nanoparticles, as the voltage increases
- SOM-4 an electron microscope image of a 2D array of AuNPs and a study of their size
- SOM-5 UV-VIS spectra of some of the AuNPs with varied ligands
- SOM-6 some additional details of modeling

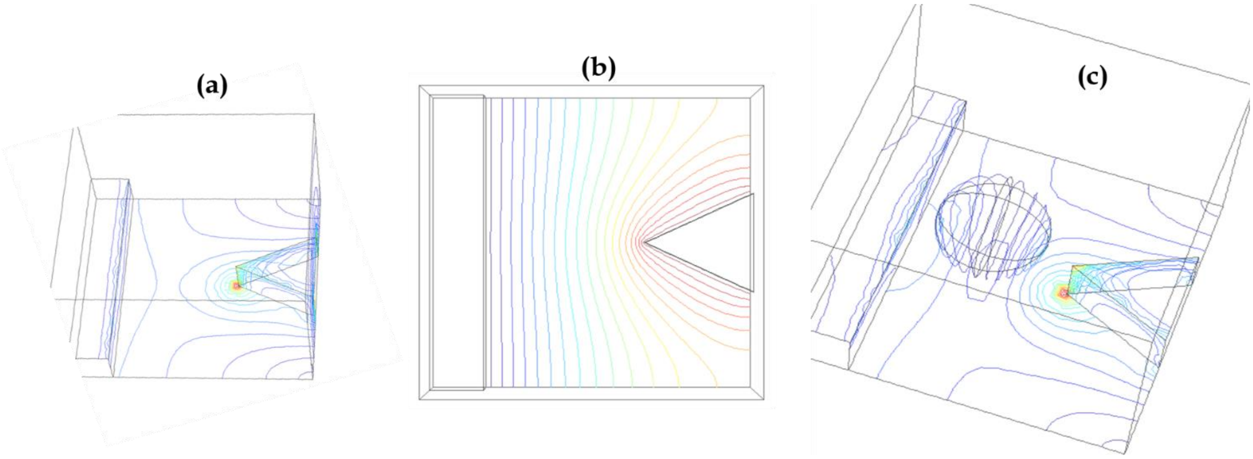


**Figure SOM-1** (a) a 3D model of electric field lines for two electrodes (a triangular prism on the right, and a long, rectangular prism on the left . (b) a top view of the electric field lines for two electrodes. (c) the repeat of the image from Fig. 1 in the main text. The results are numerically obtained

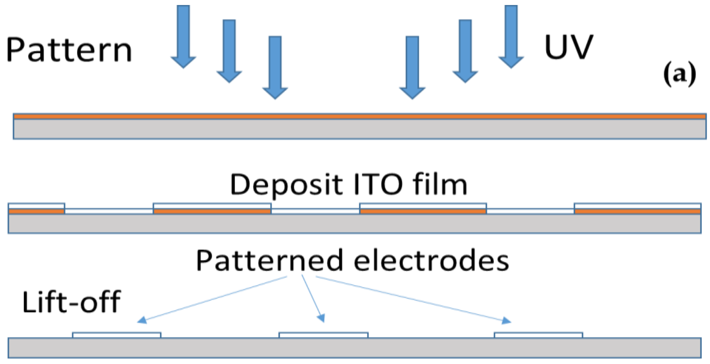


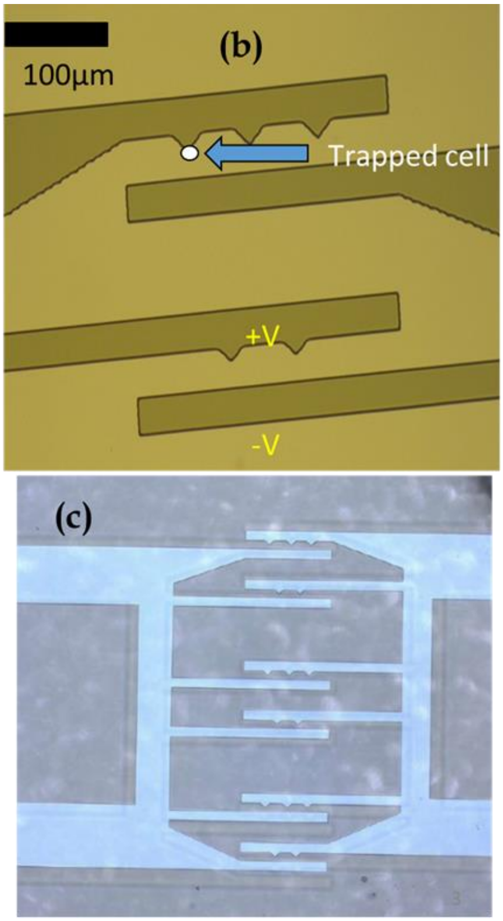


**Figure SOM-2** (a) a side view of the main steps in the microfabrication process used to prepare the electrodes on a chip; (b) a top-view image in a microscope during the microfabrication; (c) a microscope image of the finalized electrode array.

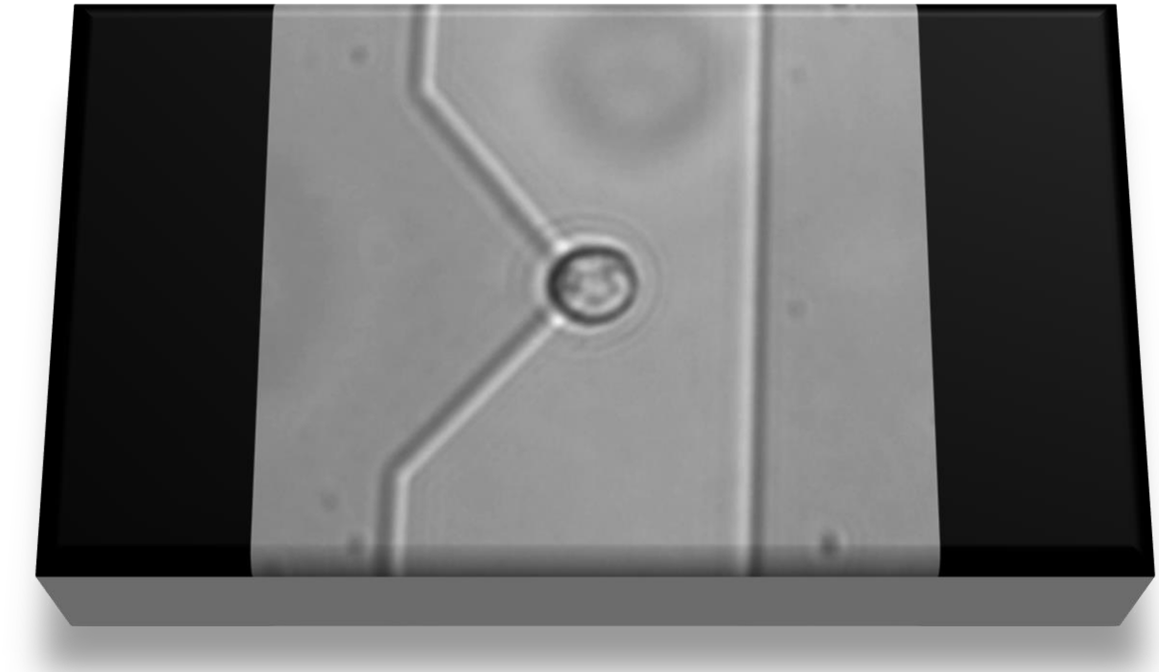

**Figure SOM-3** a movie recording of the deformation process of gold nanoparticles-enriched white blood cells; If the movie doesn't play, please e-mail the corresponding author.

**(a)**

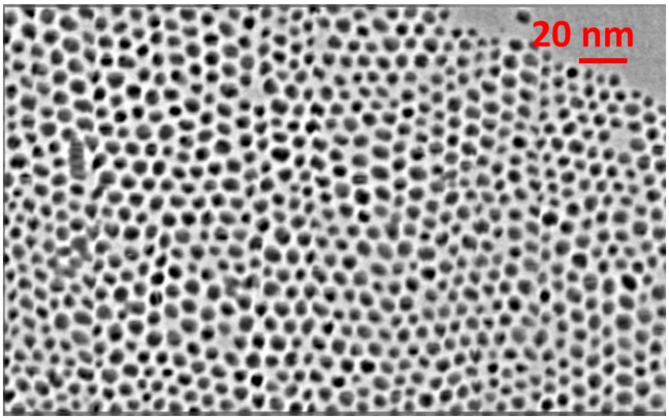


**Figure SOM-4**

(a) electron miscroscope image of a 2D layer of gold nanoparticles (AuNPs);

(b) the distribution of the sizes, raw data (hashed blue lines) and a fit (full red line).

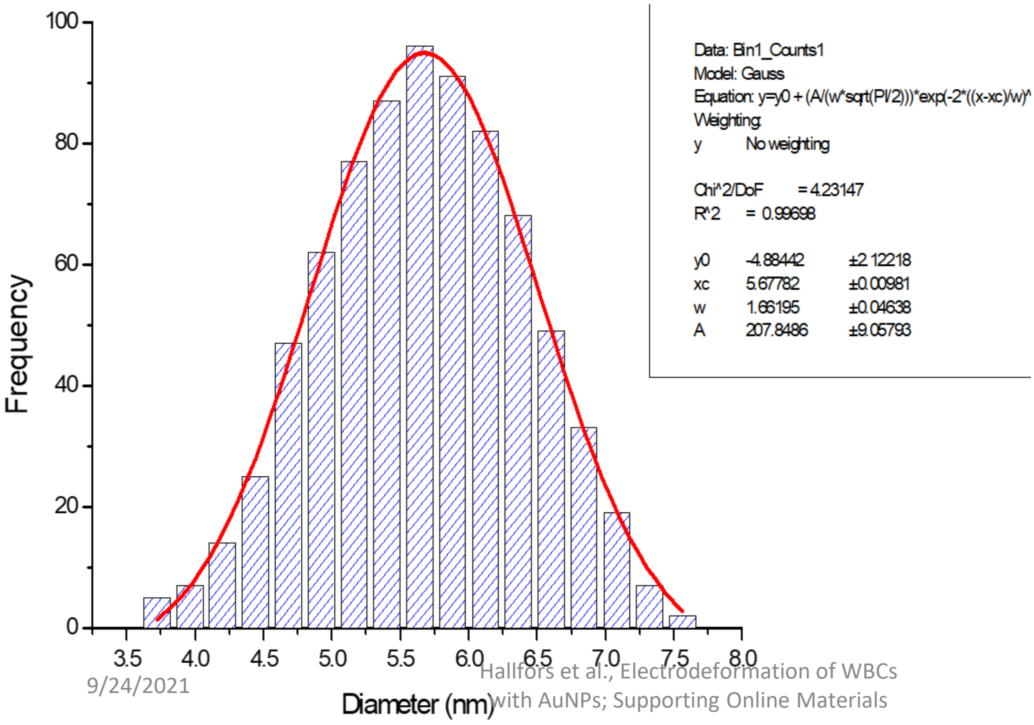

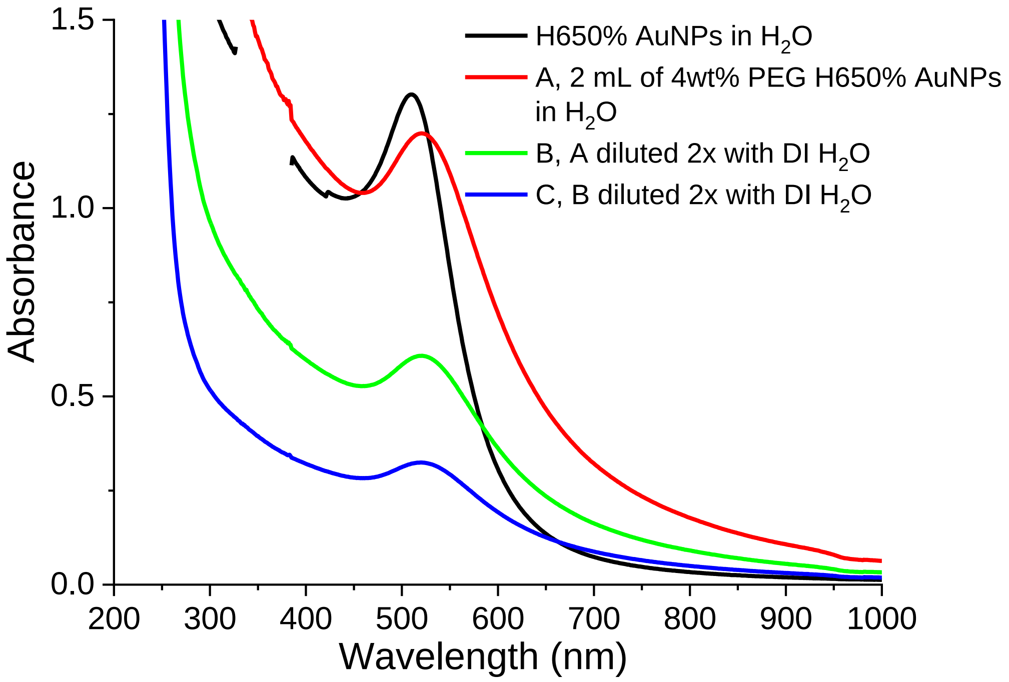


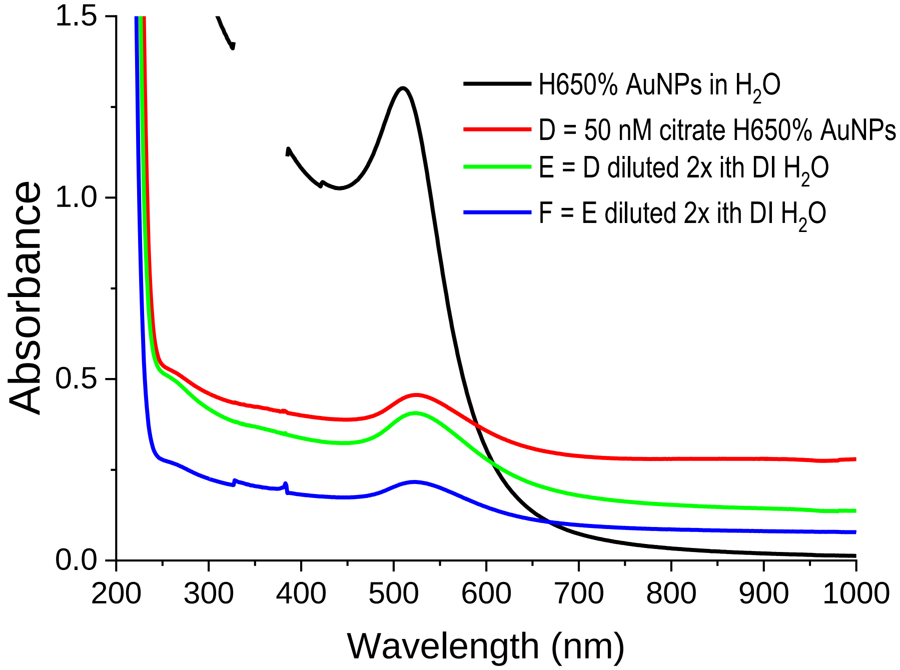


**Figure SOM-5** some main aspects of the procedure for making AuNP solutions, together with absorbance spectra

- While preparing 4wt% PEG H650% AuNPs, we have added 2 mL of H650% AuNPs to 80 mg of thiolated PEG and then vortexed the resulting mixture.
- Next, we took 1 mL of this solution and diluted with 1 mL of DI water.
- For preparing citrate solution, we have taken 9.5 g of H650% AuNPs and added 0.5g of 1M trisodium citrate solution to make total 10 g volume. Next, we took 5 mL of this 50mM citrate H650% AuNPs and mixed with 5 mL of DI water to obtain 25 mM citrate H650% AuNPs in water.

This model works under the assumption that the cell is perfectly spherical. The model then approximates the cytoplasm as a conducting sphere surrounded by a thin dielectric shell which is the cell membrane. Within the cell, the nucleoplasm is represented as a conducting sphere with a thin dielectric shell representing the nuclear envelope. Every phase of the cell is then described by specific dielectric parameters: permittivity (ε) and conductivity (σ). A MATLAB code was then written for a double-shell lymphocyte based off of general code written by Ronald Pethig [14]. Specific physical and dielectric parameters of a lymphocyte were obtained by Yulia Polevaya et al and were inserted into the code [47].

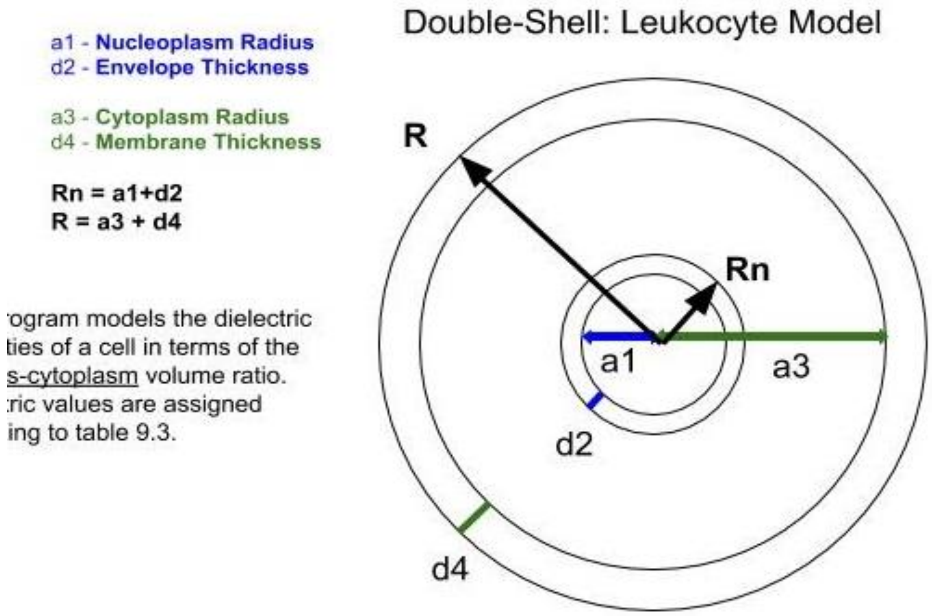


| DOUBLE SHELL MODEL **Dielectric Parameters** | | | | | Nucleoplasm (Np) | | Nuclear envelope (Ne) | | Cytoplasm (Cp) | | Membrane (M) | |
|---|---|---|---|---|---|---|---|---|---|---|---|---|
| | Np:(a1) | Ne:(d2) | Cp:(a3) | M:(d4) | **σnp**(S/m): (kc1) | **εnp**: (kp1) | **σne**(S/m): (kc2) | **εne**: (kp2) | **σcp**(S/m): (kc3) | **εcp**: (kp3) | **σm**(S/m): (kc4) | **εm**: (kp4) |
| Normal B-Cell | 3.37E-06 | 4.00E-08 | 3.93E-06 | 7.00E-09 | 2.04 | 120 | 1.11E-02 | 106 | 1.31 | 60 | 5.60E-05 | 12.8 |

**Figure SOM-6** additional details of the numerical work to model dielectric response of cells, following our upgrades of the model in Pethig, R. (2017). Dielectrophoresis: Theory, Methodology and Biological Applications. ]
